# Survey of extracellular communication of systemic and organ-specific inflammatory responses through cell free messenger RNA profiling in mice

**DOI:** 10.1101/2021.12.14.472685

**Authors:** Jiali Zhuang, Arkaitz Ibarra, Alexander Acosta, Amy P. Karns, Jonathan Aballi, Michael Nerenberg, Stephen R. Quake, Shusuke Toden

**Author notes:** Contributed equally (Jiali Zhuang, Arkaitz Ibarra). Correspondence to: Shusuke Toden, Molecular Stethoscope, Lab 360, 269 E Grand Avenue, South San Francisco, CA 94080.

## Abstract

Inflammatory and immune responses are essential and dynamic biological processes that protect the body against acute and chronic adverse stimuli. While conventional protein markers have been used to evaluate systemic inflammatory response, the immunological response to stimulation is complex and involves modulation of a large set of genes and interacting signaling pathways of innate and adaptive immune systems. Therefore, there is a need for a non-invasive tool that can comprehensively evaluate and monitor molecular dysregulations associated with inflammatory and immune responses. Here we utilized cell-free messenger RNA (cf-mRNA) RNA-Seq whole transcriptome profiling to assess lipopolysaccharide (LPS) induced and JAK inhibitor modulated inflammatory and immune responses in mouse plasma samples. Considering that, both organ-specific recruitment of immune cells and organ resident bespoke immune cells contributes to restoration of organ homeostasis, we also examined LPS-induced gene-expression dysregulation of multiple organs to shed light on organ crosstalk. Cf-mRNA profiling displayed a pattern of systemic immune responses elicited by LPS and dysregulation of associated pathways. Moreover, attenuation of several inflammatory pathways, including STAT and interferon pathways, were observed following the treatment of JAK inhibitor. Lastly, we identified the dysregulation of liver-specific transcripts in cf-mRNA which reflected changes in the gene-expression pattern in this biological compartment. Collectively, using a preclinical model, we demonstrated the potential of plasma cf-mRNA profiling for systemic and organ-specific characterization of drug-induced molecular alterations that are associated with inflammatory and immune responses.

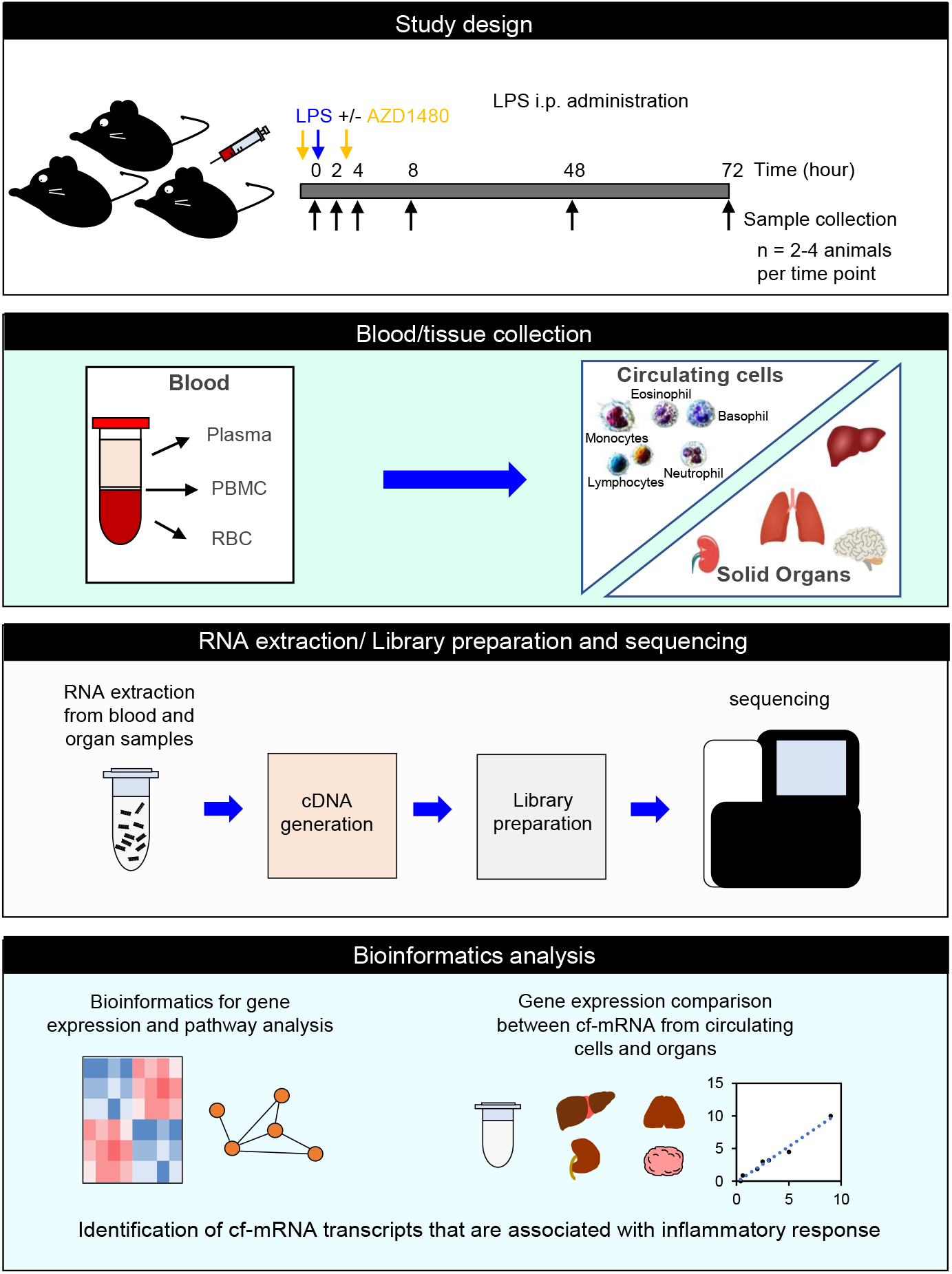

## INTRODUCTION

Inflammation is an important self-defense mechanism that is typically triggered by invading pathogens, tissue injuries, tumor growth or the onset of pathological autoimmune activities. Although inflammation is crucial for protecting the body from harmful stimuli and initiating the healing process, chronic inflammation has been recognized to contribute to multiple non-infectious diseases including type 2 diabetes, non-alcoholic fatty liver disease (NAFLD), rheumatoid arthritis, inflammatory bowel disease, cardiovascular diseases and certain types of cancer (*1*). Therefore, inflammation has become a key target for drug development in several diseases. For instance, several biologics such as infliximab, adalimumab, certolizuma pegol and golimumab are currently used to neutralize the inflammatory mediator TNF-α or to block its receptors (*2*). Moreover, currently several signaling pathways that regulate cytokine response have been targeted as a potential alternative approach to suppress inflammation (*3*). In particular, Janus kinase (JAK), a family of intracellular, non-receptor tyrosine kinases which transduce cytokine-mediated signals through the JAK-STAT pathway, has emerged as a potential therapeutic candidate for drug development (*4*–*6*). Accordingly, a number of JAK inhibitors have been approved for the treatment of rheumatoid arthritis (Baricitinib) and psoriasis (Tofacitinib), with several other drugs being tested in ongoing clinical trials (*7*, *8*). Considering the potential of immune modulators, such as JAK inhibitors, for the treatment of inflammation-related chronic diseases, there is a need for an approach that can comprehensively assess the efficacy of drugs through evaluation of target engagement and downstream target modulation.

Lipopolysaccharide (LPS) is an endotoxin derived from the outer membrane of Gramnegative bacteria which triggers a potent acute inflammatory reaction *in vivo*. LPS is known to stimulate multiple cell types including monocytes, dendritic cells, macrophages and B cells through binding to the Toll-like receptor 4 (TLR4) complex and activate downstream pathways including the IKK/NFκB pathway (*9*). The activated immune cells subsequently release cytokines and chemokines and results in further recruitment and reprograming of other immune cells, inducing systemic inflammation (*9*, *10*). Due to its well-characterized activation mechanisms, LPS has been widely used as a tool to study acute inflammation in both *in vivo* and *in vitro* models (*11*–*14*).

Although there are several existing blood-based tests, such as high-sensitivity C-reactive protein (hsCRP) and fibrinogen, that are used clinically to evaluate systemic inflammation (*15*, *16*), there are currently no non-invasive approaches that could reliably and comprehensively assess the molecular dysregulation that are associated with inflammatory responses in a hypothesis-independent manner. Emerging evidence indicates that the molecular profile of circulating cell-free messenger RNA (cf-mRNA) reflects organ-specific molecular alterations (*17*–*22*). We have developed a highly robust RNA-Seq based assay that can accurately quantify the cf-mRNA transcriptome. We also assessed transcriptional dysregulation of cf-mRNA in patients with Alzheimer’s disease (*21*), liver disease (*22*) and hematological cancers (*23*) and demonstrated the prospect of clinical utility of the cf-mRNA platform. Here, we examine the robustness of cf-mRNA profiling for the monitoring of LPS and immune modulator induced inflammatory responses using a preclinical mouse model. Our data demonstrate the potential utility of cf-mRNA profiling for the evaluation of drug-induced molecular dysregulations of systemic and organ-specific inflammatory responses and may be useful as a biomarker for drug development and clinical trials.

## MATERIALS AND METHODS

### Study design and sample collection

For all studies six to ten-week-old C57BL/6 mice were used. We conducted two independent studies. One of which is focused on the collection of blood sample following treatment of LPS and JAK inhibitor (Study 1) and the other focused on the collection of organ and blood samples following LPS insult (Study 2). For study 1, animals were treated with LPS (3mg/kg body weight: VetOne) (time = 0). LPS was administered through intraperitoneal injection. Formulations were prepared by diluting LPS in sterile saline solution, and by combining AZD1480 (a JAK2 inhibitor) (30mg/kg body weight; Thermo Fisher) in 0.5% hydroxypropyl methylcellulose + 1% Tween-80, with the pH adjusted to 3.0 for AZD1480. Blood was collected from the mice via cardiac puncture at 2, 4, 8, 24, 48 and 72 hours post LPS administration, and control animals treated with vehicle. For the administration of AZD1480, two doses of AZD1480 were administered orally, 1 hour prior to and 3 hours post LPS administration. Animals used for the study 1 are summarized in **Table 1**. For study 2, animals were treated with LPS (3mg/kg body weight: VetOne and diluted in saline) (time = 0) via intraperitoneal injection and blood and selected organs were collected 4 24, 42,78 hours post LPS administration. Blood samples were collected by cardiac puncture and selected organs including lung, brain, kidney and liver were harvested from exsanguinated mice. The organs were dissected half longitudinally and one section was placed in trizol and the other section flash frozen on dry ice. All samples were stored at −80 °C. For each treatment group 2-4 biological replicates were evaluated (**Table 2**). All animals were housed, and experiments were performed by BTS Research, San Diego, CA. All housing, handling and treatment protocols were approved and handled according to the standards of the Institutional Animal Care and Use Committee.

**Table 1:** Blood collection and animal allocation summary (Study 1)

| Time points | Plasma |  |  | PBMC |  |  | RBC |  |  |
| --- | --- | --- | --- | --- | --- | --- | --- | --- | --- |
|  | Untreated | LPS | LPS + AZD | Untreated | LPS | LPS + AZD | Untreated | LPS | LPS + AZD |
| 0h | 4 |  |  | 5 |  |  | 5 |  |  |
| 2h |  | 5 | 6 |  | 3 | 4 |  | 3 | 4 |
| 4h |  | 3 | 6 |  | 3 | 4 |  | 3 | 4 |
| 8h |  | 3 | 4 |  | 3 | 2 |  | 2 | 3 |
| 48h |  | 3 |  |  | 3 |  |  | 3 |  |
| 72h |  | 3 |  |  | 3 |  |  | 3 |  |

Note: numbers represent samples collected at each time point.

**Table 2:** Blood and tissue sample collection summary (Study 2)

| Time | Blood (Plasma) |  | Liver |  | Lung |  | Brain |  | Kidney |  |
| --- | --- | --- | --- | --- | --- | --- | --- | --- | --- | --- |
| Time points | Untreated | LPS | Untreated | LPS | Untreated | LPS | Untreated | LPS | Untreated | LPS |
| 0h | 3 |  | 3 |  | 3 |  | 3 |  | 3 |  |
| 4h |  | 5 |  | 5 |  | 3 |  | 2 |  | 3 |
| 24h |  | 3 |  | 3 |  | 3 |  | 3 |  | 3 |
| 48h |  | 4 |  | 4 |  | 3 |  | 2 |  | 3 |
| 72h |  | 3 |  | 2 |  | 3 |  | 2 |  | 3 |

Note: numbers represent samples collected at each time point.

### RNA extraction, library preparation and whole-transcriptome RNA-seq

RNA was extracted from up to 500 μl of plasma using QIA amp Circulating Nucleic Acid Kit (Qiagen) and eluted in 15 μl volume. ERCC RNA Spike-In Mix (Thermo Fisher Scientific, Cat. # 4456740) was added to RNA as an exogenous spike-in control according to manufacturer’s instructions. Agilent RNA 6000 Pico chip (Agilent Technologies, Cat. # 5067-1513) was used to assess the integrity of extracted RNA. RNA samples were converted into a sequencing library as described previously (*23*). Qualitative and quantitative analysis of the NGS library preparation process was conducted using a chip-based electrophoresis and libraries were quantified using a qPCR-based quantification kit (Roche, Cat. # KK4824). Sequencing was performed using Illumina NextSeq500 platform (Illumina Inc), using paired-end sequencing, 76-cycle sequencing. Base-calling was performed on an Illumina BaseSpace platform (Illumina Inc), using the FASTQ Generation Application. For sequencing data analysis, adaptor sequences were removed and low-quality bases were trimmed using cutadapt (v1.11). Reads shorter than 15 base-pairs were excluded from subsequent analysis. Read sequences greater than 15 base-pairs were aligned to the mouse reference genome GRCm38 using STAR (v2.5.2b) with GENCODE vM14 gene models. Duplicated reads were removed using the samtools (v1.3.1) rmdup command. Gene expression levels (in the unit of transcripts per million (TPM)) were calculated from de-duplicated BAM files using RSEM (v1.3.0).

### Identification of non-peripheral blood cell and tissue-specific cf-mRNA transcripts

In plasma, a large portion of cell-free transcripts remain peripheral blood cell origin (*24*). These transcripts that are derived from peripheral blood cells are in general originated from the bone marrow, but not from organs of interest. Therefore, in order to identify which transcripts are enriched in plasma, but not in peripheral blood cells, we compared the transcriptional profiles of plasma and compared to that of peripheral blood mononuclear cells (PBMC). To identify peripheral blood cell (PBC) transcripts, the following criterion was used. We define R_ij_ as the ratio of the TPM in plasma over the TPM in PBMC fraction for gene *i* in mouse *j*. Similarly, we define r_ij_ as the ratio of the TPM in plasma over the TPM in RBC fraction for gene *i* in mouse *j*. Let L_i_ denote the number of plasma samples where gene *i* is detected (TPM > 3), N_i_ denotes the number of plasma samples where R_ij_ > 3, and M_i_ denotes the number of plasma samples where r_ij_ > 3. Gene *i* is considered as “non PBC”‘ if L_i_ >= 6 AND M_i_ > L_i_ * 0.9 AND N_i_ > L_i_ * 0.9. Tissue (cell-type) specific transcripts are defined as transcripts whose expression in a particular tissue (cell-type) is > 5-fold higher than all the other tissue types (cell-types). Tissue (cell-type) transcriptome expression levels were obtained from the following two datasets: BodyMap for gene expression across 17 human tissues and Immgen for gene expression in endothelia cell, epithelial cell and fibroblastic reticular cell. In this study we only considered the tissue/cell-type specificity for non-PBC transcripts.

### Differential expression analysis and pathway enrichment analysis

Differential expression analysis was implemented with DESeq2 (v1.12.4) (*25*) using read counts as input. Genes with fewer than 5 total reads across the entire cohort were excluded from subsequent analysis. Benjamin-Hochberg correction was used to correct for multiple testing and obtain adjusted p-values. Pathway enrichment analysis was conducted using Ingenuity Pathway Analysis (IPA) software version 47547484 or Gene Ontology (R package “topGO”). The complete list of differentially expressed transcripts was uploaded to IPA and Expression Analysis was used to determine pathways that are highly enriched. IPA categories including Canonical pathways and “Top diseases and bio functions” were examined.

### Computational transcriptome deconvolution analysis using non-negative matrix factorization (NMF)

Normalization was first implemented whereby the expression level of each gene in each sample was divided by the gene’s maximum value across the samples. This step is designed to rescale the expression levels among different genes to avoid a small number of highly expressed genes dominating the decomposition process. The normalized expression matrix was then subjected to NMF decomposition using sklearn.decomposition.NMF within the Python library Scikit-learn (https://scikit-learn.org/stable/). NMF decomposition achieves a more parsimonious representation of the data by decomposing the expression matrix into the product of two matrices X = WH. X is the expression matrix with n rows (n samples) and m columns (m genes); W is the coefficient matrix with n rows (n samples) and p columns (p components); H is the loading matrix with p rows (p components) and m columns (m genes). W is in a sense a summarization of the original matrix H with reduced number of dimensions. H contains information about how much each gene contribute to the components. Biological interpretation of the derived components was achieved by performing pathway analysis on the top genes that contribute the most to each component.

### Statistical analysis

Pearson’s correlation was used to examine the correlation between different bioanalytes. Within a condition/treatment group, a median of all the animals in the group was taken before correlation or fold change calculation. Student’s t-test was used to evaluate the difference between the two groups. The Benjamin-Hochberg method was used to correct for multiple testing. All statistical analyses were performed using R (3.4.4, R Development Core Team, https://cran.r-project.org/) unless otherwise stated.

## RESULTS

### Identification of functionally relevant LPS associated cf-mRNA components

LPS stimulation is a widely utilized preclinical tool to induce an acute inflammatory response and the molecular signatures of the response have been well characterized in mice (*11*–*13*). Therefore, we first investigated whether the cf-mRNA profile is altered in response to acute LPS-induced stimulation. A single dose of LPS (6 mg/kg) was administrated to each C57bl6 mice intraperitoneally. Animals were sacrificed at 2, 4, 8, 48 and 72 hours following the LPS treatment and blood and organ samples were collected (Figure 1A and Table 1). RNA was subsequently isolated from the plasma and sequenced. The evaluation of gene-expression pattern in cf-mRNA showed induction of cytokine transcripts, as early as 2 hours after the LPS treatment, followed by a subsequent regression of cytokine transcripts 8 hours after the treatment (Supplementary Figure 1). We then compared the cf-mRNA transcriptome of untreated and LPS treated animals (4-hour post LSP treatment) and identified 750 differentially expressed cf-mRNA transcripts (476 upregulated and 274 downregulated, FDR < 0.05 was used as the cut-off criterion). We chose 4-hour post-treatment samples for the comparison as cytokine transcripts appear to be at highest levels at this time point in the plasma compared to other time points. We performed canonical pathway analysis using IPA (Qiagen) using transcripts that were upregulated following LPS treatment. Pathway analysis identified several key inflammation-associated pathways including, acute phase response (*p* < 0.0001), interferon signaling (*p* < 0.0001), IL-6 signaling (*p* < 0.0001) and TLR signaling (*p* < 0.0001) as the most enriched pathways (Figure 1B). Furthermore, the most significant common upstream regulator identified by IPA was LPS (*p* < 0.0001), followed by major LPS response mediators such as Interferon-γ, Stat1, TNF and Toll-like receptor 3 (Figure 1C). Collectively, these data indicate that prominent inflammatory signals that are typically dysregulated by LPS stimulation are detected in the plasma cf-mRNA of mice treated with LPS.

**Figure 1:**
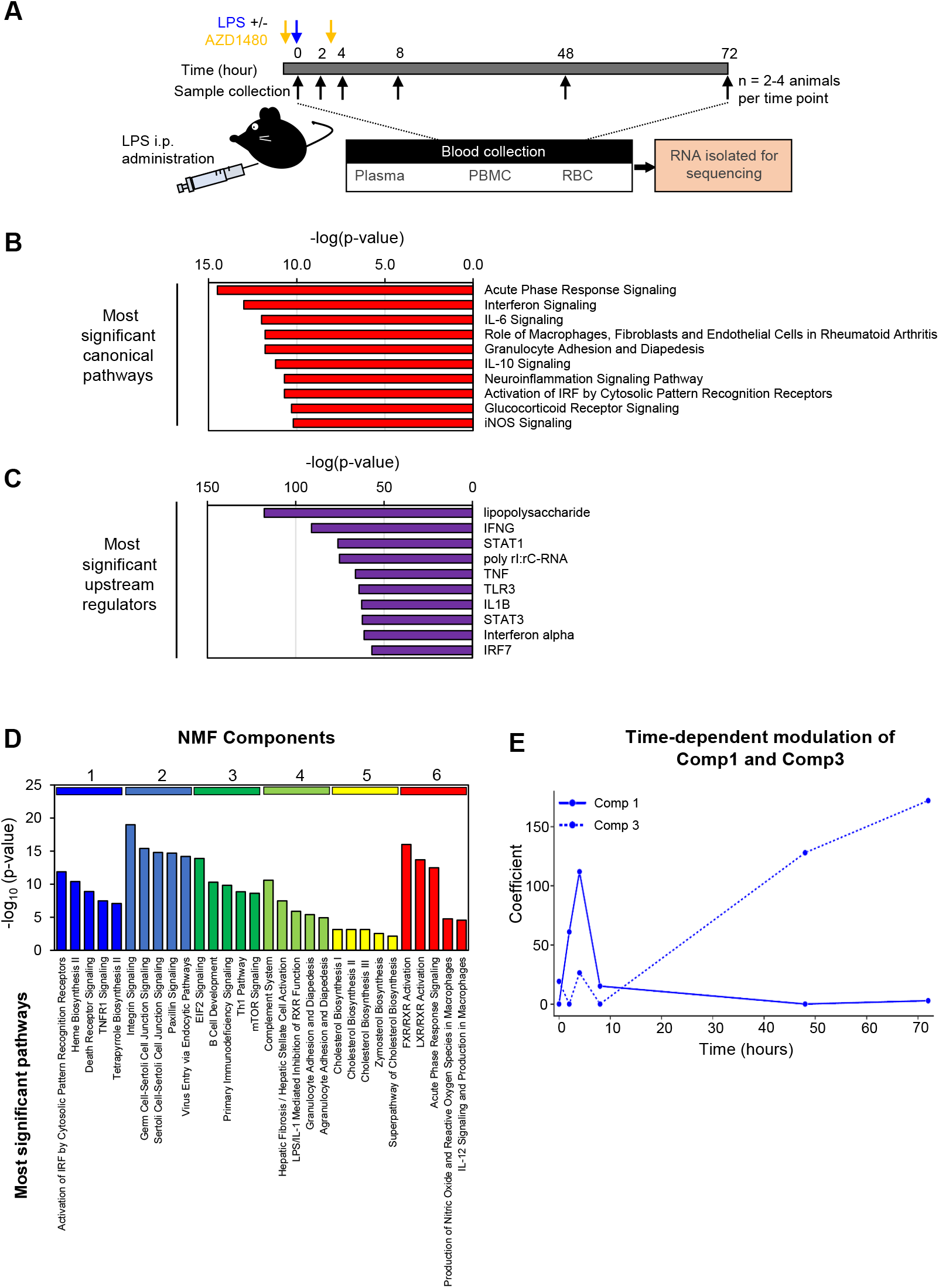
Identification of 6 cf-mRNA sub-clusters following LPS treatment. (**A**) A schematic overview of the experimental design. (**B**) Most significant IPA canonical pathways identified using 750 dysregulated cf-mRNA transcripts as inputs (Control vs 4 h. post LPS treatment). (**C**) Upstream regulators identified using 750 dysregulated cf-mRNA transcripts as inputs (Control vs 4 h. post LPS treatment). (**C**) Most significant canonical pathways for individual NMF components. (**D**) Temporal patterns of component 1 (solid line) and component 3 (dashed line) transcripts.

Next, we performed unsupervised clustering of cf-mRNA transcripts using Non-negative matrix factorization (NMF) to evaluate whether these transcripts share expression patterns across various time points and conditions and form gene clusters. NMF analysis resulted in identification of six clusters of gene components (Supplementary Figure 3). IPA pathway analysis of the individual clusters showed that these components are enriched in distinct biological processes and pathways (Figure 1D). For example, Component 1 transcripts are enriched in biological processes such as “innate immune response” (*p* < 0.0001) and “response to cytokines” (*p* < 0.0001) (Figure 1D), while Component 3 transcripts are enriched in lymphocyte-specific markers and lymphocytic -related pathways (Figure 1D). We then evaluated the temporal patterns of expression level changes for each component following the LPS stimulation. The expression levels of Component 1 transcripts showed substantial increase following LPS treatment with the highest expression level observed at 4-hours post LPS time point (Figure 1E). Subsequently, the Component 1 transcripts declined and returned to the basal levels by 48 hours. The response we observed for this cluster is consistent to that of cytokine response. In contrast, Component 3 transcripts were unaffected during the early stages of the LPS stimulation but increased substantially at 48-hour post LPS treatment time point, indicative of a delayed response of lymphocytic lineage cells (Figure 1E and Supplementary Figure 2). This observation is also consistent with previous reports that an LPS-insult leads to enhanced survival and hence the accumulation of lymphocytes (*26*, *27*). Our results demonstrated that LPS-induced transcriptional changes in cf-mRNA are consistent with previously known cellular and molecular events triggered by LPS and highlights the ability of the assay to comprehensively monitor molecular alterations in cf-mRNA that are associated with inflammatory response.

### Cf-mRNA profiling captures anti-inflammatory property of immunomodulators

Given the increasing focus on immuno- and inflammatory modulations as a potential drug development strategy, we next evaluated whether cf-mRNA profiling can be used to monitor the effects of immune modulators. To assess the efficacy of immune modulators on LPS-induced transcriptional changes, we treated mice with AZD1480, orally twice, 1 hour prior and 2 hours after LPS treatment. AZD1480 is a JAK inhibitor which is known to preferentially inhibits JAK2. The AZD treatment substantially attenuated the LPS-induced elevation of the Component 1 transcripts (Figure 2A), which was enriched for immune response and cytokine response pathways. The comparison between LPS treated samples with or without AZD resulted in identification of 87 transcripts that are significantly down-regulated in the cf-mRNA of AZD treated group at 2-hour post LPS time point. IPA upstream regulator analysis revealed that these dysregulated transcripts were enriched in downstream targets of interferon cytokines (Figure 2B). Transcription factor STAT1, a key mediator of JAK signaling, ranked as the most enriched upstream regulator, indicating that AZD indeed inhibited the JAK/STAT pathway. When examining the median fold change of the downstream target genes of STAT1 and the interferons relative to the untreated control, AZD significantly attenuated the expression levels of these transcripts (Figure 2C). Furthermore, several notable interferon-γ signaling pathway-associated transcripts that were upregulated by LPS (USP18, IFIT1, CXCL10, OASL1, IFIT3B and IFIT3) were suppressed by AZD (Figure 2D). Collectively, these data indicate that the transcriptional plasma cf-mRNA profile appears to reflect the anti-inflammatory effects of JAK inhibitor and suggest that the cell-free transcriptome can be used to monitor molecular regulation of immunomodulators.

**Figure 2:**
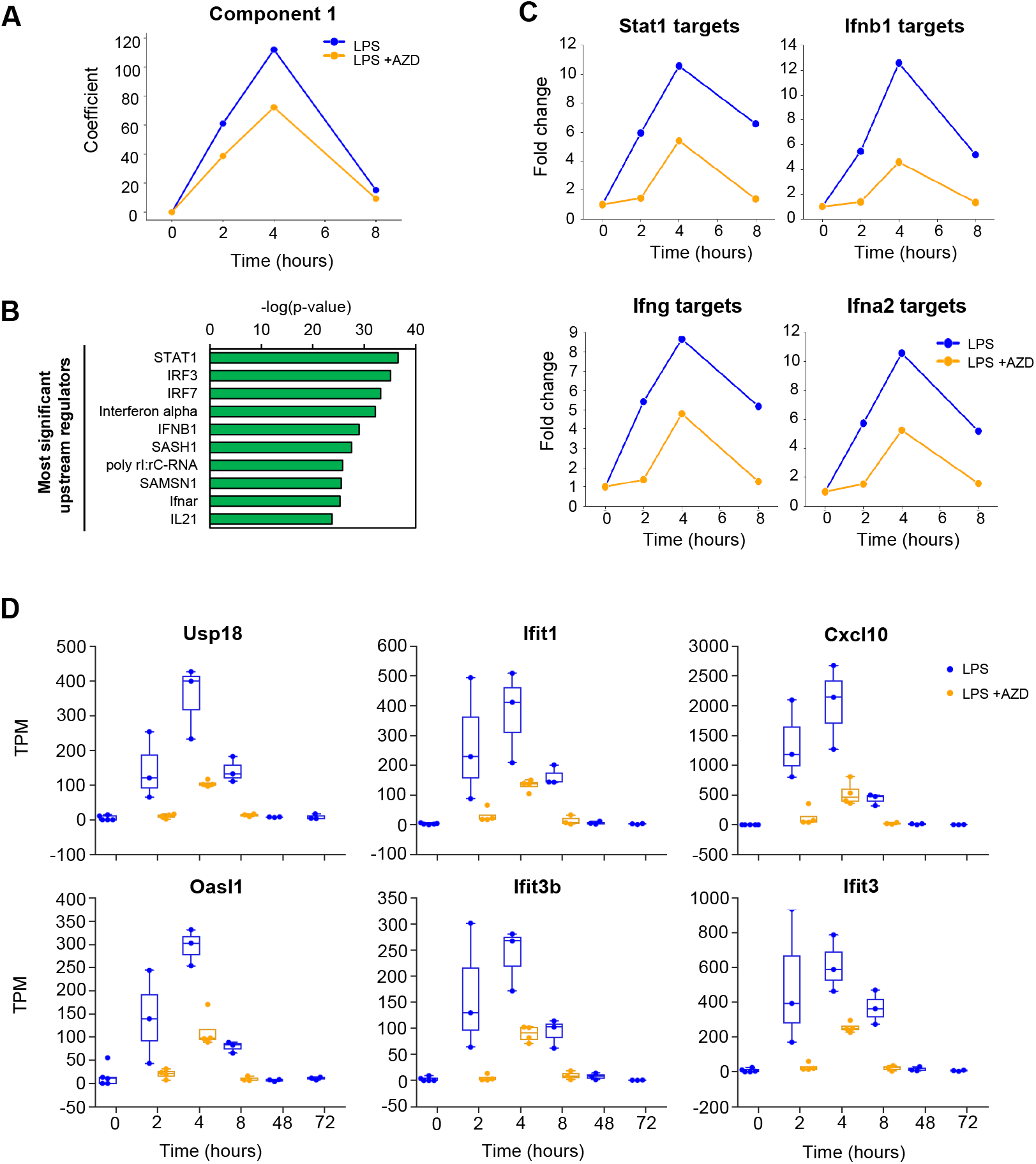
Cf-mRNA profiling captures anti-inflammatory property of immunomodulators. (**A**) Time-dependent temporal patterns of component 1 transcripts following LPS treatment with (blue) or without (orange) AZD. (**B**) Upstream regulators identified using 87 dysregulated cf-mRNA transcripts as inputs (2 h. post LPS treatment with or without AZD). (**C**) Average fold changes of target transcripts of JAK/STAT related upstream regulators relative to the untreated controls. (**D**) Temporal changes in expression levels of Interferon-γ related transcripts.

### Identification of tissue-specific gene-expression signals for inflammatory responses

We previously reported that both circulating blood cells and resident tissue cells contribute to the composition of cf-mRNA transcriptome in humans (*23*). After demonstrating that cf-mRNA profiling can be used to monitor the LPS-induced inflammatory and immune responses, we next investigated whether transcriptional alterations that are occurring in the solid tissues following LPS stimulation can be detected in cf-mRNA. First, we evaluated the transcriptome profiles of matched mouse plasma, PBMCs and red blood cells by separating different blood fractions using Ficoll-Paque and conducted RNA sequencing on the each fraction. The comparison of geneexpression profiles between these fractions resulted in identification of 1,054 transcripts that are highly enriched in the plasma cf-mRNA fraction relative to both the PMBC and RBC fractions. We termed these transcripts “non-peripheral blood cell (PBC) transcripts”, since these gene transcripts are enriched in the cell-free fraction of the blood when compared to the peripheral blood cells. Next, we investigated the expression levels of these non-PBC transcripts across various organs and cell types that are published in well-established public datasets (BodyMap & Immgen). A portion of the non-PBC transcripts that we have identified in the cf-mRNA were indeed organ/tissue specific (Figure 3A). In particular, the quantification of organ-specific transcripts in cf-mRNA suggests that the majority of the organ-specific transcripts in the circulation appear to be derived from liver, brain, heart and muscle (Figure 3B).

**Figure 3:**
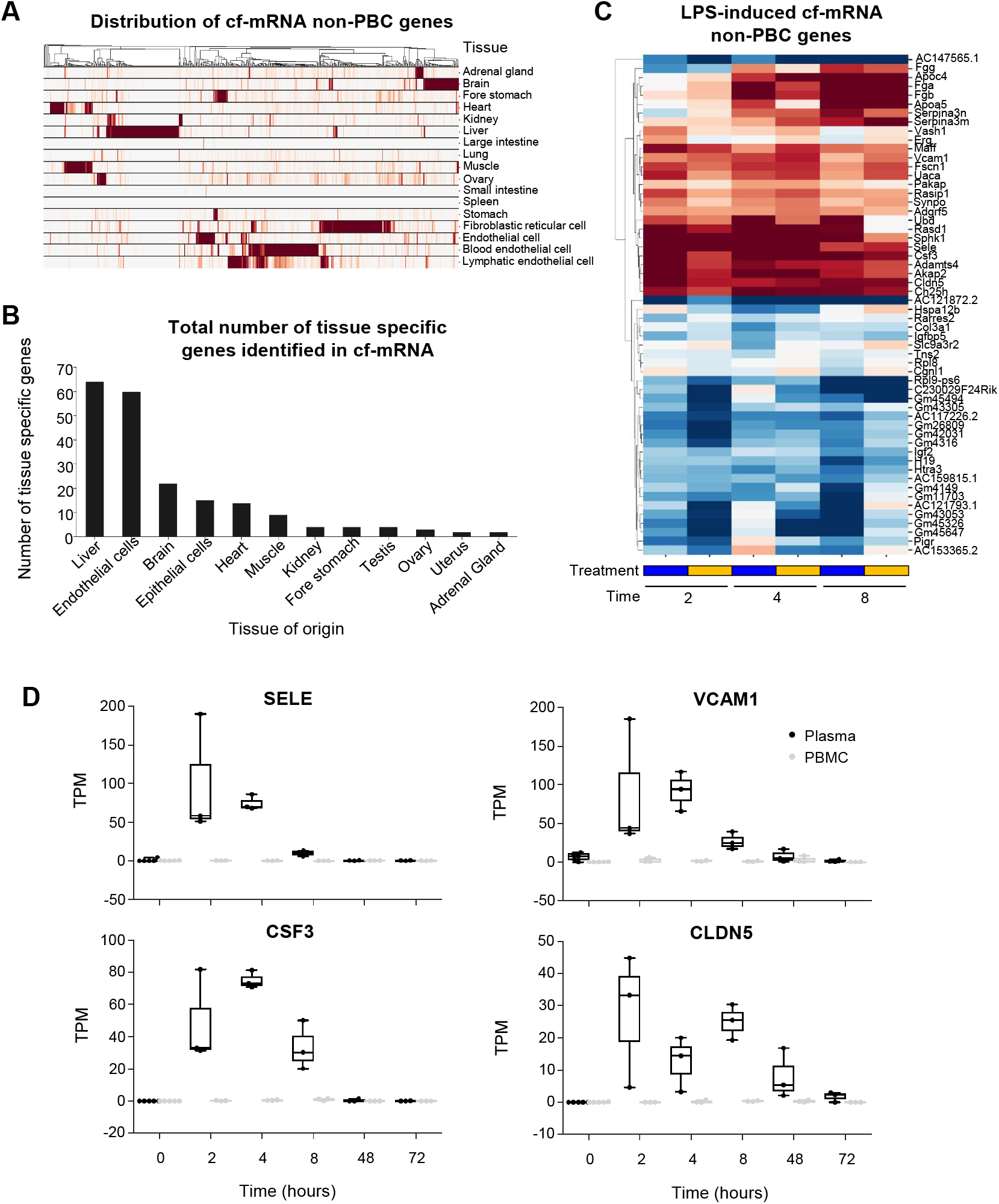
Identification of tissue-specific gene-expression signals for inflammatory responses. (**A**) Expression patterns of non-peripheral blood cell (PBC) transcripts across tissues and cell types. Each row represents a tissue or cell type while each column represents a gene. Each column (gene) is normalized by its maximum value. A hierarchical clustering was performed on the columns. (**B**) Number of tissue specific transcripts. A transcript with expression level in a particular tissue >5 fold higher than any other tissue is considered specific to that particular tissue. (**C**) A heatmap depicting fold changes relative to baseline at various time points for non-PBC transcripts. Only transcripts that are significantly differentially expressed compared to baseline (FDR < 0.01) and with TPM > 30 in at least one condition are shown. Hierarchical clustering was performed on the transcripts (rows). Animals treated with LPS treatment with (blue) or without (orange) AZD. (**D**) Examples of LPS-induced expression changes for non-PBC transcripts. Expression levels in the plasma samples were denoted in black while those in the matching PBMC samples were denoted in grey.

Out of 1054 non-PBC transcripts that we have identified, 167 transcripts were significantly dysregulated following LPS stimulation (FDR < 0.05). Most of the organ-specific transcripts dysregulated by LPS treatment were liver-specific (71%). These transcripts were significantly elevated in 4- and 8-hour post LPS treatment time points (Supplementary Figure 4). A collection of the most significantly differentially expressed transcripts (FDR < 0.01 with TPM > 30 in at least one condition) are shown in Figure 3C. In particular, vascular cell adhesion molecule (VCAM1) and E-selectin (SELE) showed a marked increase in plasma following LPS stimulation (Figure 3D). These two genes are known to encode cell surface adhesion molecules that play crucial roles in the adhering of circulating leukocytes to vascular endothelial cells and their subsequent extravasation to the site of inflammation (*28*). Both genes have been shown to be up-regulated following LPS treatment in human endothelial cells through an NF-κB dependent mechanism (*28*, *29*). Furthermore, granulocyte colony-stimulating factor (CSF3) is known to encode a cytokine that stimulates hematopoiesis of the phagocytic neutrophils and its precursors and play an essential part in innate immune response (*30*). The levels of CSF3 increased significantly following LPS induction and returned to the basal level by 48-hour post-treatment time point (Figure 3D). Moreover, we showed that LPS stimulation increased the expression level of Claudin 5 (CLDN5) in the circulation, a gene which encodes tight junction protein that is highly specific to endothelial cells. In addition, CLDN4 has been shown to be a key regulator of endothelium permeability (*31*); our data were consistent with a previous study (*32*) (Figure 3D). Upregulation of these transcripts were only observed in plasma, not PBMC, suggesting that these tissue-specific transcripts are highly enriched in the plasma portion of blood. Collectively these data showed that cf-mRNA geneexpression profiles appear to reflect inflammatory processes in the tissues and could potentially be used for organ-specific assessment of inflammatory responses.

### LPS induced transcriptional changes in the organs are reflected in the cf-mRNA fraction

To investigate whether LPS-induced transcriptional alternations in specific organs are reflected in cf-mRNA, we examined gene-expression profiles of multiple organs following LPS stimulation. We conducted an additional experiment where we collected both plasma corresponding tissue samples (liver, kidney, lung and brain tissues) from LPS treated mice 4, 24, 48 and 72 hours post-LPS treatment (Figure 4A). Differentially expressed transcripts were identified using the untreated animals as the reference and pathway enrichment analysis was performed on differentially expressed transcripts using IPA. In the organs we examined, transcripts up-regulated 4 hours after LPS stimulation were significantly enriched in “acute phase response”, “IL-6 signaling” and “Interferon signaling” pathways, suggesting that LPS-induced inflammatory effects were observed. Indeed, in all the organs analyzed, LPS was the most prominent upstream regulator identified by IPA, along with other well-known inflammation regulators such as Interferons and STAT1. Furthermore, we assessed the time-dependent dynamics of the LPS-induced response by evaluating the average fold-changes of the downstream target genes of the inflammation regulators relative to untreated controls (Figure 4B). The analysis showed elevation of LPS-related immune pathways in the tested tissue types at 4-hour post treatment time point followed by steady attenuation (Figure 4B). Consistently, the time-dependent dynamics of genes involved in the major inflammatory pathways followed similar patterns (Figure 4C).

**Figure 4:**
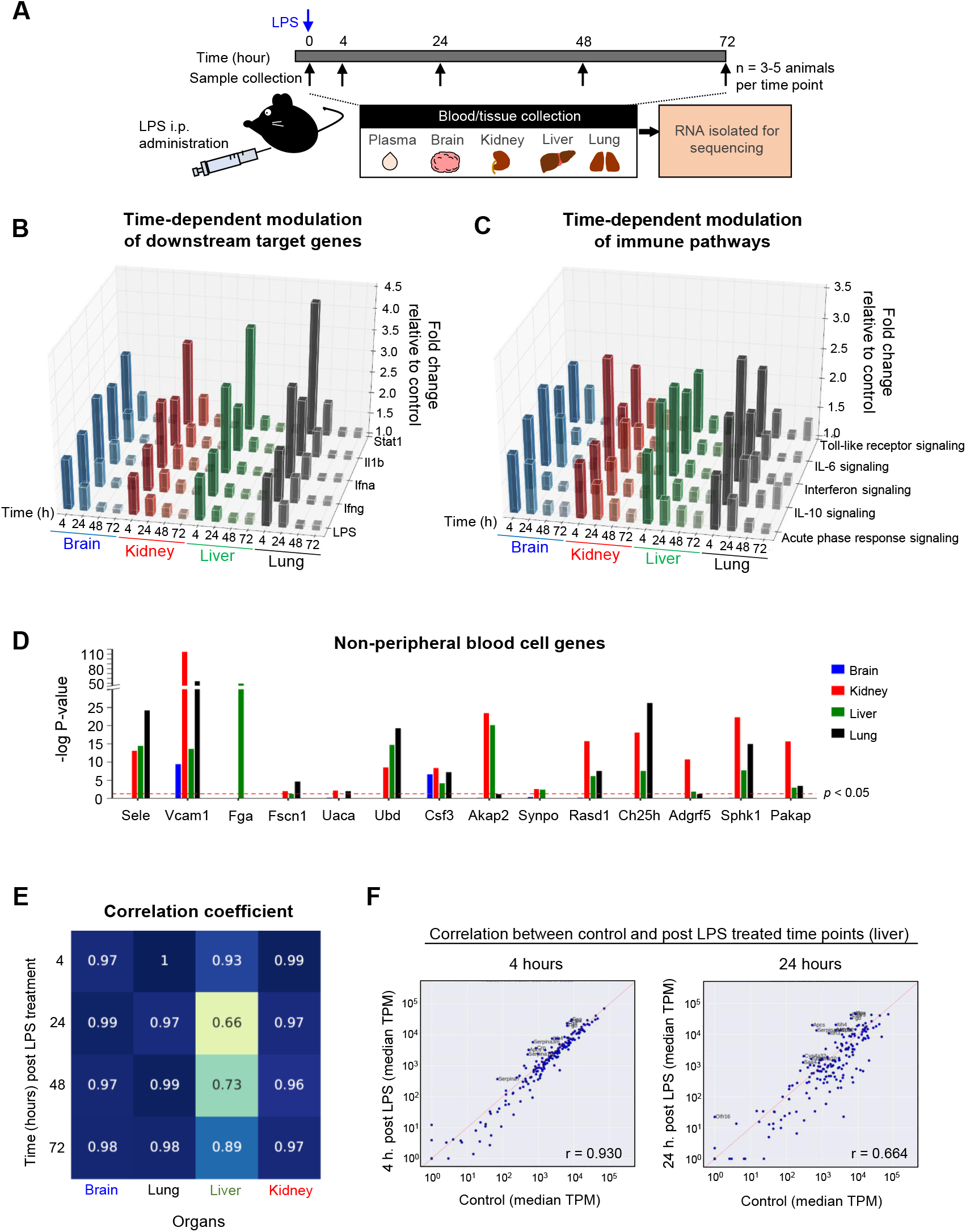
Identification of organ-specific transcript dysregulation. (**A**) A schematic overview of the experimental design. (**B**) Bar plots showing the median fold changes of transcripts in inflammation related canonical pathways relative to untreated controls. (**C**) Bar plots showing the median fold changes of downstream target transcripts for immune related regulators Stat1, Il1b, Interferon-α, Interferon-β and LPS relative to untreated controls. (**D**) Bar plots showing the statistical significance (corrected p-value) of differential expression in the tissues 4 hours after LPS treatment. Those transcripts were chosen because they were significantly upregulated in the plasma and with TPM > 30. The red dashed line indicates the p = 0.05 significance level. (**E**) Heatmap showing the correlation of tissue specific transcript expression levels between LPS-treated samples and untreated control samples. The columns correspond to different tissues while rows correspond to different time points after treatment. The numbers in each grid represents the Pearson Correlation Coefficient for each comparison. (**F**) Scatter plots comparing expression levels of liver specific transcripts 4 hours (left) and 24 hours (right) after LPS treatment against untreated controls (x-axis).

In the previous section, we identified a number of cf-mRNA transcripts not expressed in peripheral blood cells that were dysregulated following LPS treatment. Considering that transcripts not expressed in peripheral blood cells are likely to be originated from solid tissues, we examined whether similar gene-expression dysregulation can be observed in the transcriptomes of solid tissues. We focused specifically on the transcripts not expressed in peripheral blood cells that displayed increased levels of cf-mRNA transcripts at 4-hour post LPS treatment time point, in order to conduct a direct comparison between cf-mRNA and tissue profiles at a matching time point. We showed that all of tissue-specific transcripts were significantly up-regulated in at least one of the studied organs, with most of transcripts being significant in multiple organs (Figure 4D).

Next, we investigated the LPS-induced molecular dysregulation in organs and examined transcriptional alterations of organ-specific transcripts. Transcriptional profiles of post-LPS stimulation time points were compared to that of untreated control. In the lung, brain and kidney, the expression levels of organ-specific transcripts remain relatively unchanged at all time points, indicating that LPS treatment had minimal influence on transcriptional dysregulation on organ-specific genes in lung, brain and kidney (Figure 4E and Supplementary Figure 5). However, in the liver tissue we observed a substantial alteration of organ-specific transcripts, with the highest dysregulation observed at 24-hour post LPS. Of the dysregulated organ-specific transcripts, we identified several transcripts that were categorized in the “acute phase response” pathway (IPA) including complement system components (C3), haptoglobin (HP), fibrinogens (FGA, FGB and FGG) and serum amyloid A proteins (SAA1, SAA2, SAA3, SAA4) (Supplementary Figure 6). Dysregulation of these organ-specific genes in the liver tissues are consistent with previous reports (*33*, *34*). Collectively, these data indicate that a subset of organ-specific transcripts in the liver appear to be dysregulated following LPS-stimulation.

### cf-mRNA profiling captures liver-specific LPS-induced inflammatory response

Considering that we identified dysregulation of several organ-specific transcripts in the liver following LPS stimulation, we further examined time-dependent gene-expression alterations of liver-specific transcripts in cf-mRNA following LPS stimulation. Since, study 1 has more data within the first 8 hours for animals that were treated with LPS, we examined cf-mRNA profiles of animals that were treated with LPS in study 1. In general, the majority of liver-specific transcripts increased at 4-hour time point and were further elevated at 8-hour time point and this pattern was reflected in the median cf-mRNA liver-specific transcripts (Figure 5A). Concordantly, using a counting threshold of TPM > 5, the total number of detectable liver-specific transcripts significantly increased at 4- and 8-hour post-LPS treatment time points compared to the 2 hour-time point (Figure 5B). These data indicate that liver-specific genes are modulated by LPS-stimulation.

**Figure 5:**
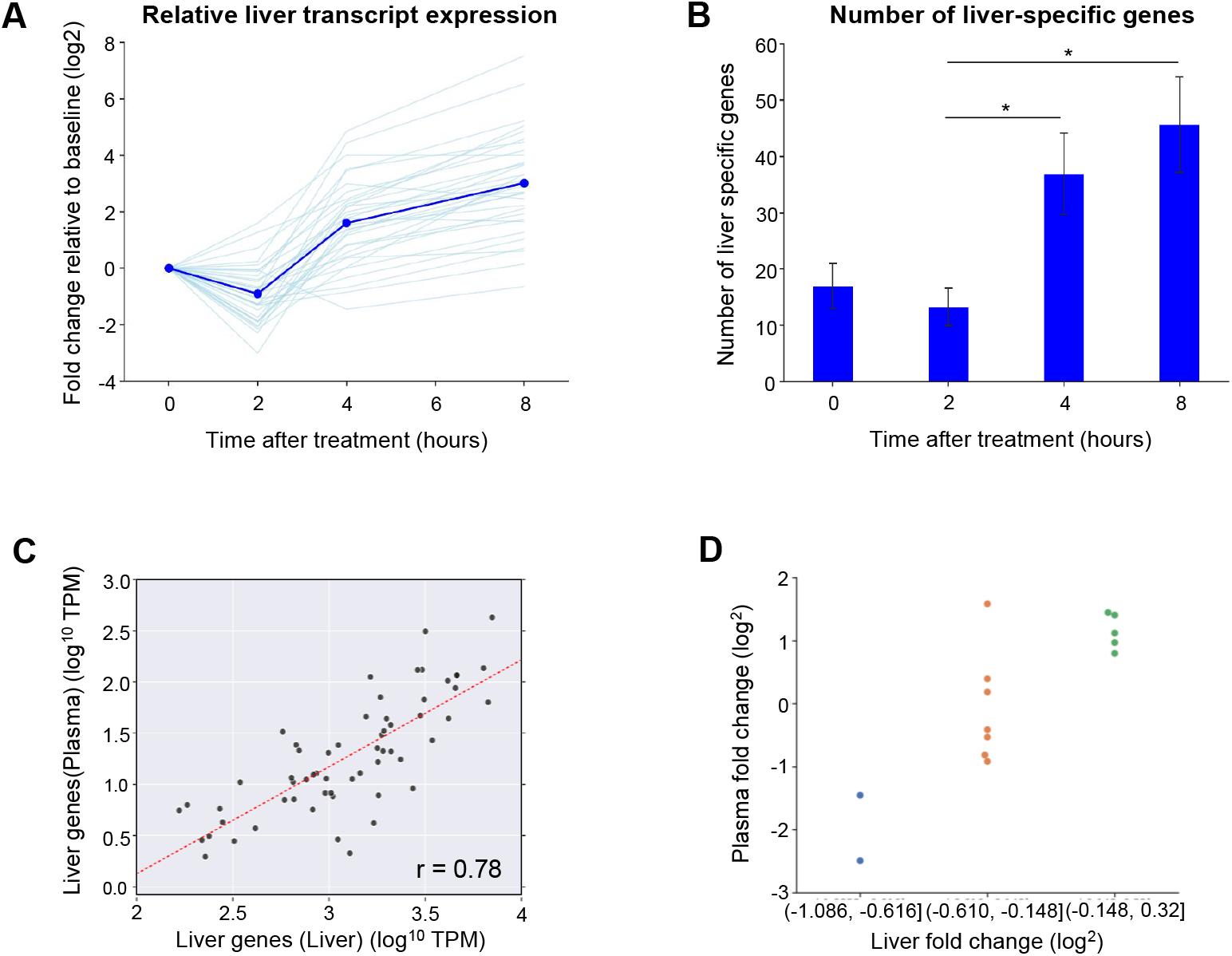
LPS-induced transcriptional changes of liver-specific transcripts are reflected in cf-mRNA. (**A**) Gene-expression changes of liver specific transcripts following LPS treatment (represented as fold change). Each light blue line represents one liver specific transcript, the blue curve represents the median of all the liver specific transcripts. The fold changes are relative to untreated controls. (**B**) Number of liver specific transcripts detected in plasma samples at specific time points (TPM > 5 was used as a detection cut off for individual genes). (**C**) Scatter plot directly comparing expression levels of liver specific transcripts in the liver tissue (x-axis) and the plasma sample (y-axis). The comparison shown was based on liver tissues and plasma samples harvested from mice 8 hours after LPS treatment. (**D**) Liver specific transcripts were grouped by their expression fold change in the liver tissue between 4-hour and 8-hour after LPS treatment. The group with higher fold change in the liver tissue also have higher fold change in the plasma and vice versa.

In order to compare the liver-specific gene expression profiles between liver tissue and plasma cf-mRNA, we conducted a correlation analysis of the TPM levels of liver-specific transcripts in the plasma and tissue at 8-hour post-LPS treatment. This time point was chosen for the analysis due to the highest number of liver-specific transcripts were detected in plasma cf-mRNA at this particular time point. We identified a significant linear correlation between the expression levels of liver-specific transcripts in cf-mRNA and that of liver tissue (Pearson’s correlation: *r* = 0.78 and *p* < 0.0001) (Figure 5C), indicating that the transcriptional dysregulation of liver-specific transcripts in the liver appear to be reflected in the cf-mRNA profile. Furthermore, we examined relative foldchange of the individual liver-specific transcripts between the untreated time point and time points 4 and 8 hours post-LPS stimulation and evaluated the corresponding abundance of individual transcripts between liver tissue and plasma as a fold-change. As expected, the liver-specific transcripts with higher fold-change in the liver had correspondingly higher fold-change in the plasma (Figure 4E). Accordingly, there was a linear correlation between the fold-changes calculated from liver tissue and plasma (Pearson’s correlation, *r* = 0.82, *p* < 0.001, Supplementary Figure 7). Taken together, these data suggest that liver-specific transcripts in the plasma compartment could be used to evaluate the molecular response in the liver to a therapeutic agent.

## DISCUSSION

Inflammatory and immune responses are complex biological responses involving dynamic modulation of a large set of genes and pathways of the innate and adaptive immune systems. In the present study, we performed RNA-Seq of cf-mRNA to characterize LPS-induced systemic and organ-specific inflammatory responses using a mouse model. We demonstrated that cf-mRNA RNA-sequencing is able to provide comprehensive insights of LPS-induced systemic and organ-specific inflammation as well as the modulatory effects of a JAK inhibitor. Furthermore, the comparison between different blood fractions resulted in the identification of transcripts that are enriched in plasma, but not in peripheral blood cells and many of these transcripts appear to be dysregulated by LPS-stimulation. Finally, we discovered several unique liver-specific transcripts in cf-mRNA that reflected molecular alterations in the liver.

In the present study, LPS stimulation was used to evaluate inflammatory response associated molecular dysregulation in cf-mRNA. We specifically chose LPS stimulation in a mouse model since molecular characteristics of LPS stimulation have been extensively studied and well-characterized in multiple organs (*11*–*13*). While there are existing blood-based tests to evaluate the inflammatory response such as C-Reactive Protein (CRP) and cytokine panels (*35*, *36*), a small number of proteins in the circulation is not sufficient to comprehensively assess a complex dynamic process which involves modulation of multiple signaling pathways. Although there are proteomics techniques including mass spectrometry and aptamer-based multiplexing platforms that allow simultaneous quantification of a limited number of proteins in the circulation (*37*, *38*), cf-mRNA profiling using RNA-sequencing technology offers an alternative approach to robustly quantify gene expression alterations of the entire transcriptome in a hypothesis-independent manner (*21*, *23*). The opportunity to discern allelic imbalance and splice variant isoforms because of the sequencing detection strategy casts an even broader net for discovery. Furthermore, we evaluated whether cf-mRNA sequencing can be used to assess the efficacy and target engagement of therapeutic agents using this *in vivo* model. Accordingly, we examined the effects of LPS stimulation as well as anti-inflammatory effects of JAK inhibitor to suppress LPS-induced inflammation. Our data showed that a JAK inhibitor substantially suppressed inflammation associated cf-mRNA transcripts that were upregulated by LPS-stimulation. Our study is consistent with and extends the findings of a recent study where inflammatory response was suppressed by JAK inhibitor in the hepatocyte following LPS stimulation (*39*). Our approach of simultaneous monitoring of multiple transcripts in the circulation ideally positioned for drugs and drug combinations that target multiple genes or pathways. Collectively, our data demonstrated that the dysregulation of cf-mRNA in the plasma reflected the efficacy of the immune modulating drugs that are administered to the animals.

Many chronic diseases such as inflammatory bowel disease, non-alcoholic fatty liver disease (NAFLD), diabetes, rheumatoid arthritis Alzheimer’s Disease and lupus are characterized by chronic inflammation of specific organs/tissues. Unfortunately, blood-based inflammatory biomarkers alone are generally unable to quantify organ-specific immune-related molecular alterations. We and others have shown previously that a portion of messenger RNAs are known to be specific or enriched in distinct organs (*18*, *40*, *41*). Therefore, we looked for tissue-specific transcripts that are dysregulated by LPS-stimulation. Our data from both cf-mRNA and tissue transcriptome profiling indicated that the liver is the organ most affected by the LPS treatment. A direct comparison between cf-mRNA and the matching liver tissue profiles indicated that a small portion of liver-specific transcripts in cf-mRNA appear to recapitulate LPS-induced molecular dysregulation in the liver. Although we have demonstrated the significant overlap of dysregulated genes and pathways between plasma cf-mRNA and the corresponding organs in our previous clinical studies (*21*, *22*), these studies used molecular tissue profiles of independent studies and not matched tissues. In contrast, in this study we generated plasma cf-mRNA and tissue sequencing data from matched samples using a preclinical model. Our study further supports that tissue-specific molecular dysregulations are reflected in the gene-expression profile of cf-mRNA. In addition, the comparison of transcriptomic profiles between cf-mRNA and PBMC showed that the tissue-specific transcripts are detectable in plasma, but not PBMC, confirming that the tissue-derived transcripts are predominantly enriched in cell-free portion of blood and not PBMC.

In summary, we used a mouse preclinical model to demonstrate the utility of cf-mRNA for comprehensive systemic and organ-specific transcriptomic profiling of LPS induced and JAK inhibitor modulated inflammatory and immune responses. Prior studies have reported that cells release extracellular vesicles containing biologically active nucleic acids and subsequently alters the function of the recipient cells (*42*, *43*). Therefore, cf-mRNA may to some extent reflect interception of intercellular communiques. Our data highlight the potential of cf-mRNA profiling for the evaluation of drug efficacy and safety and may inform various steps of drug development and clinical trials including target engagement, pharmacodynamic monitoring, pharmacokinetic assessment and toxicological examination.

