## Supplementary Figures for "Survey of extracellular communication of systemic and organ-specific inflammatory responses through cell free messenger RNA profiling in mice"

#### Supplementary Figure 1

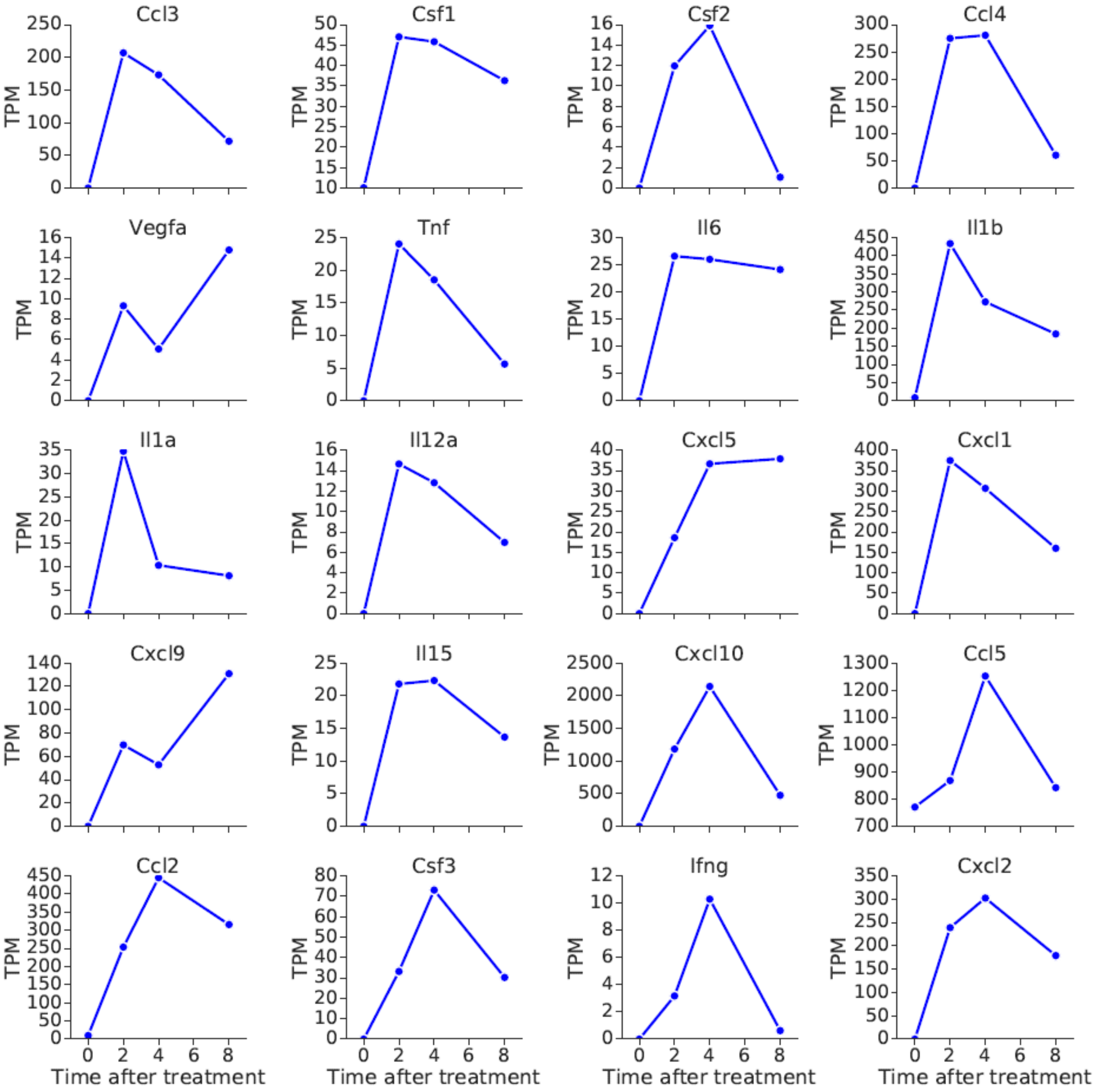


**Supplementary Figure 1:** Cytokine transcripts levels in plasma following LPS stimulation (LPS stimulation occurred at 0 hour)

### Supplementary Figure 2

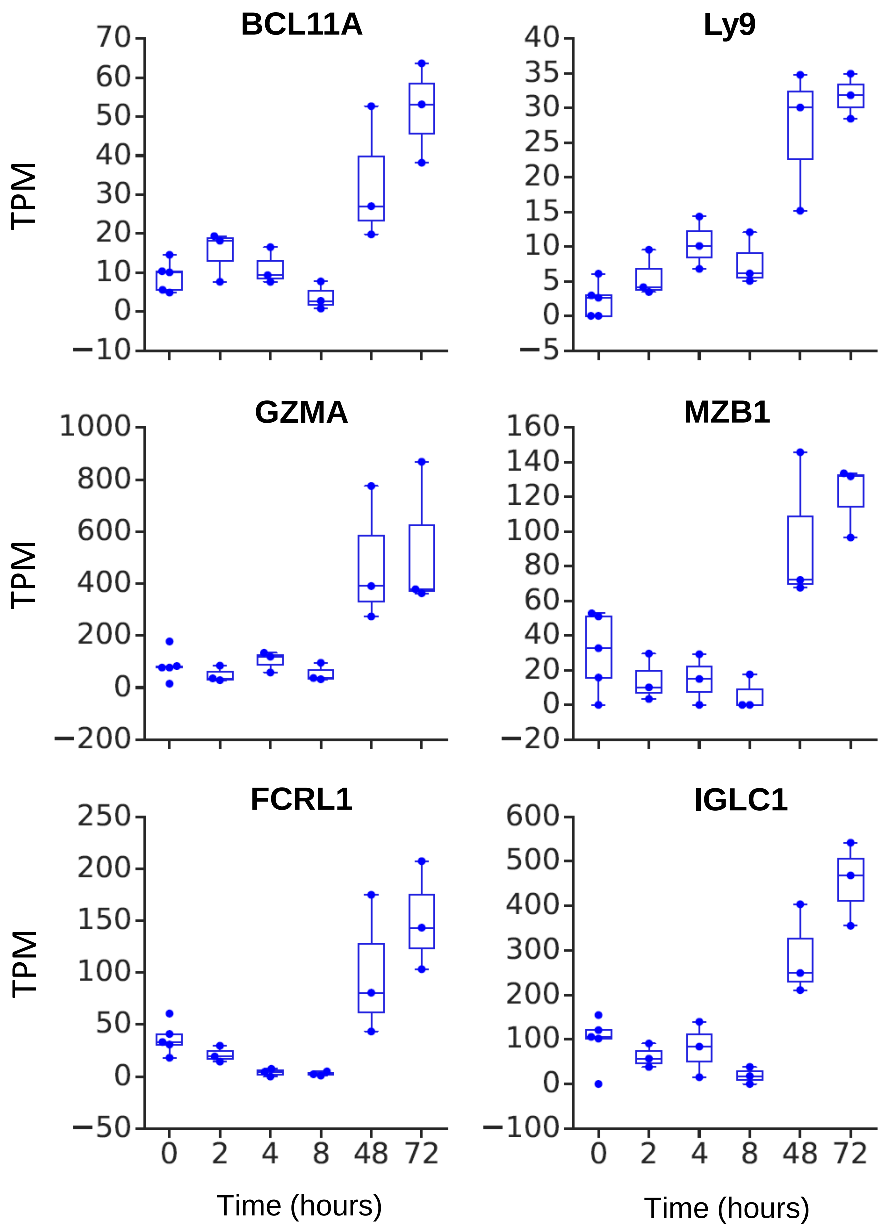


**Supplementary Figure 2:** Lymphocytic specific cf-mRNA transcripts levels in plasma following LPS stimulation

### Supplementary Figure 3

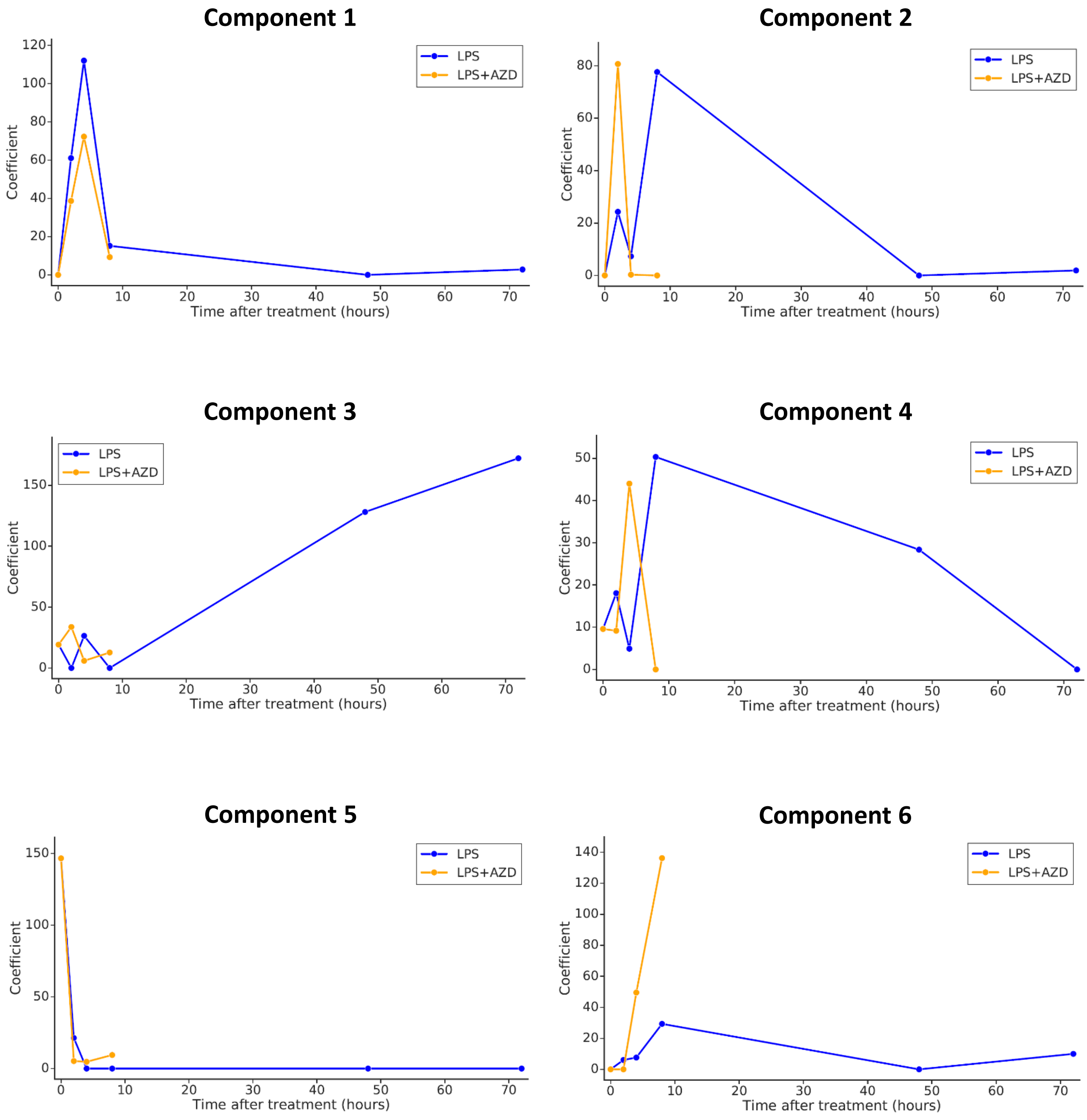


**Supplementary Figure 3:** Temporal patterns for individual NMF components for animal treated with or without AZD following LPS stimulation

### Supplementary Figure 4

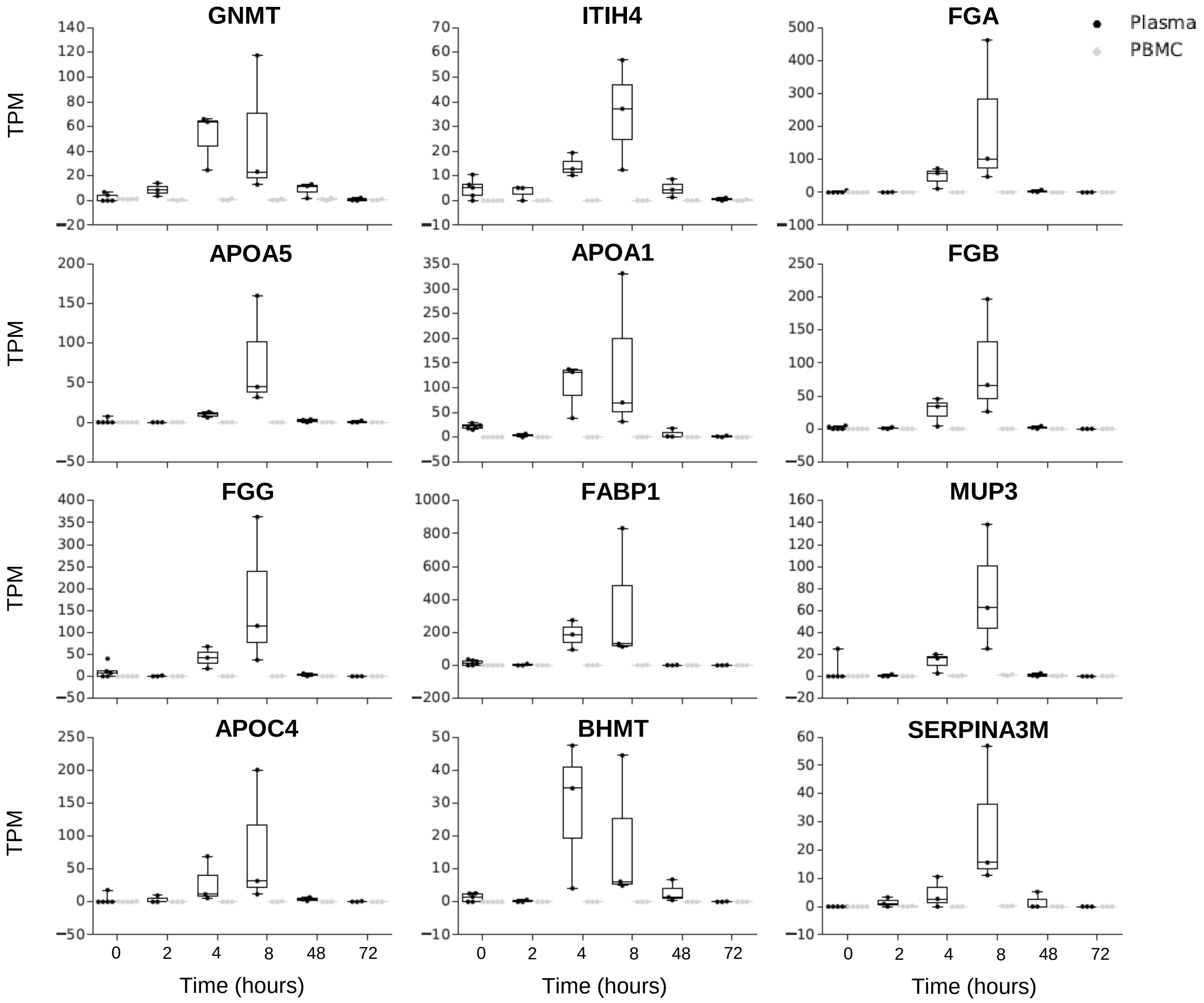


**Supplementary Figure 4:** Liver specific cf-mRNA transcripts significantly up-regulated in plasma after LPS stimulation

### Supplementary Figure 5

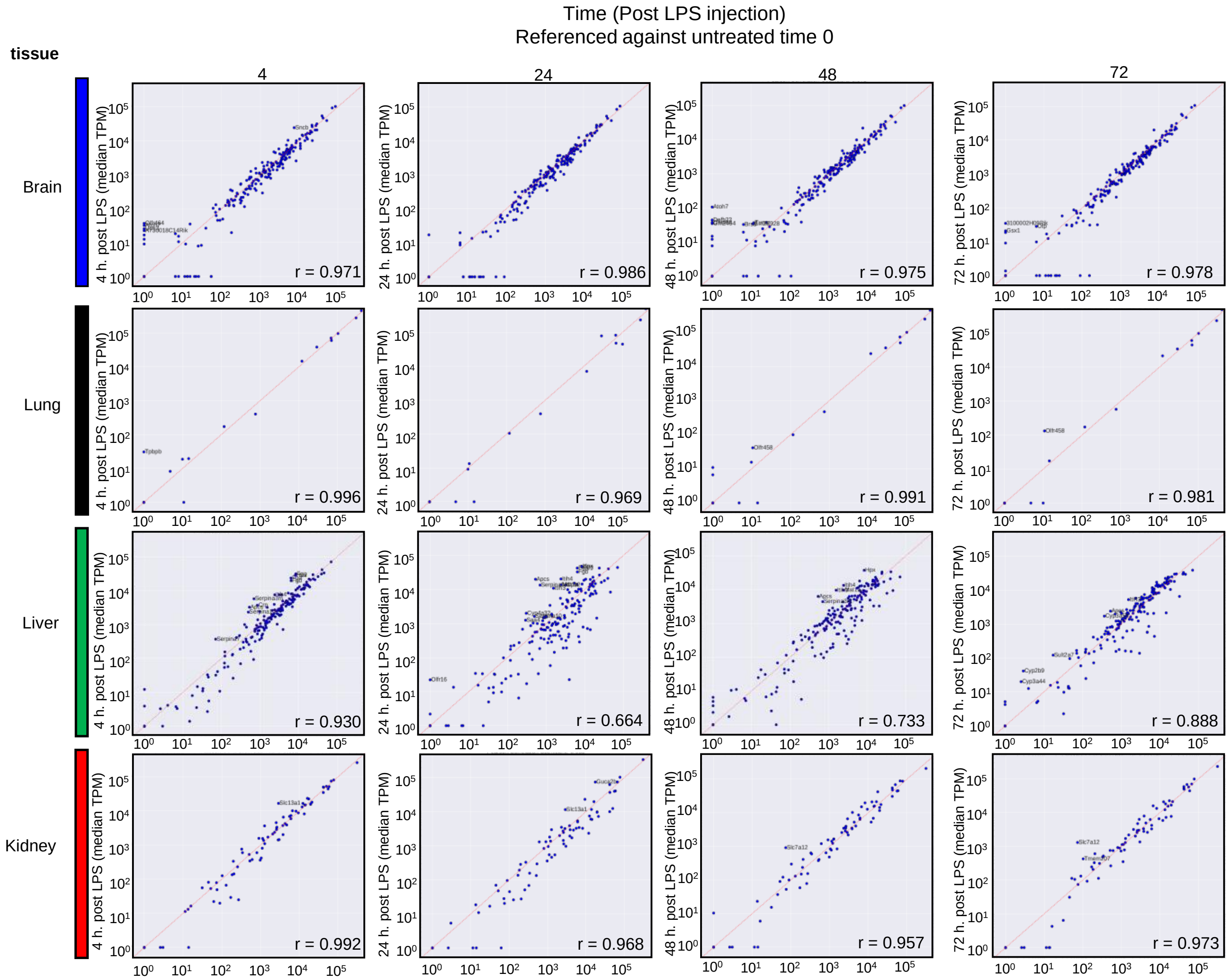


**Supplementary Figure 5:** Scatter plots showing the correlation of tissue specific transcript expression levels between LPS-treated samples and untreated control samples. In the grid rows correspond to different tissues while columns correspond to different time points after treatment. The untreated samples are always depicted in the x-axis.

### Supplementary Figure 6

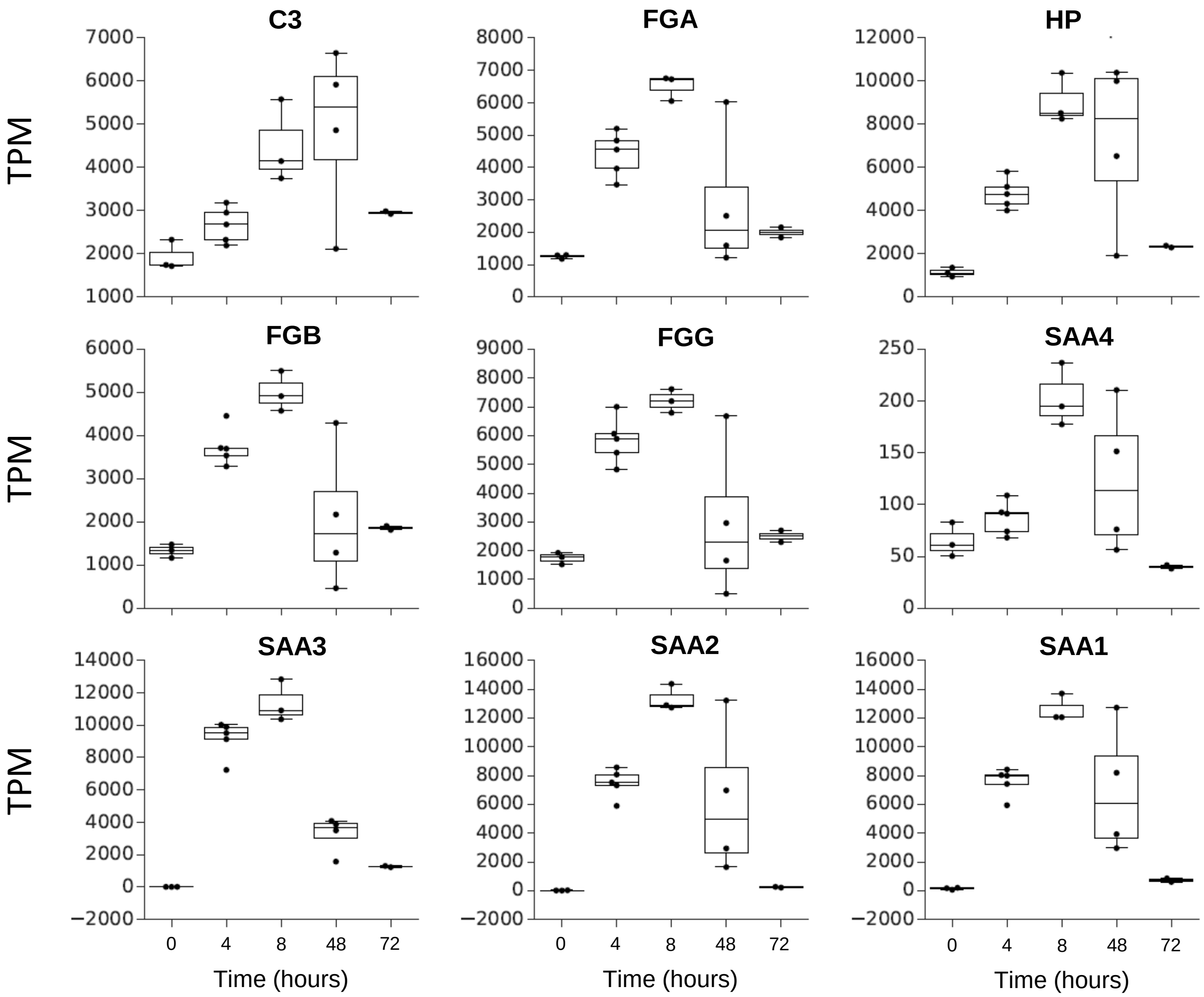


**Supplementary Figure 6:** Acute phase response transcripts levels in the liver tissue after LPS stimulation

### Supplementary Figure 7

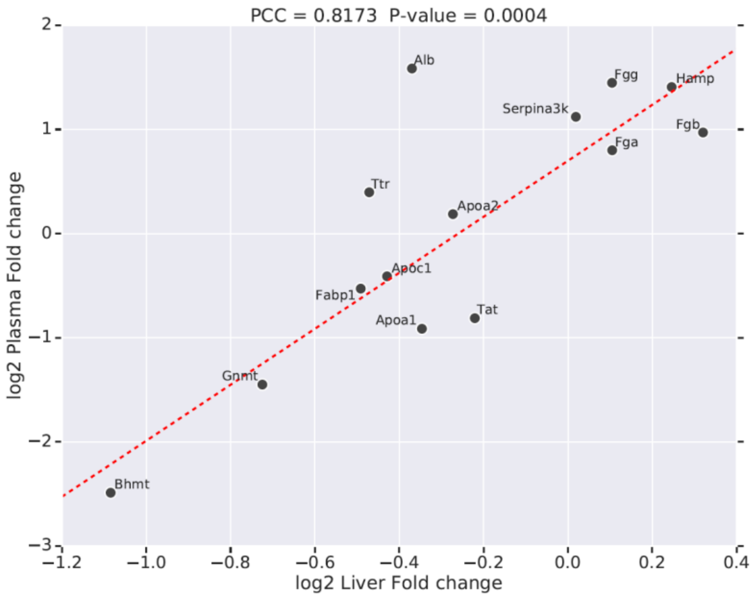


**Supplementary Figure 7:** Scatter plot showing the correlation between fold changes of liver specific transcripts (8h after LPS vs 4h after LPS) in the liver tissue (x-axis) and plasma cf-mRNA (y-axis)
